# Molecular diversity and connectivity of accessory olfactory system neurons

**DOI:** 10.1101/2022.11.08.515541

**Authors:** Nandkishore Prakash, Heidi Y Matos, Sonia Sebaoui, Luke Tsai, Tuyen Tran, Adejimi Aromolaran, Isabella Atrachji, Nya Campbell, Meredith Goodrich, David Hernandez-Pineda, Maria Herrero, Tsutomu Hirata, Julieta Lischinsky, Wendolin Martinez, Shisui Torii, Satoshi Yamashita, Katie Sokolowski, Shigeyuki Esumi, Yuka Imamura Kawasawa, Kazue Hashimoto-Torii, Kevin S Jones, Joshua G Corbin

**Author notes:** Correspondence: Joshua G Corbin, Center for Neuroscience Research, Children’s Research Institute, Children’s National Hospital, Washington DC, USA. Equal contribution (Nandkishore Prakash, Heidi Y Matos, Sonia Sebaoui).

## Abstract

Olfaction is the primary sensory modality by which most vertebrate species interpret environmental cues for appropriate behavioral outputs. The olfactory system is subdivided into main (MOS) and accessory (AOS) components which process volatile and non-volatile cues. While much is known regarding the molecular diversity of neurons that comprise the MOS, less is known about the AOS. Here, focusing on the AOS which is largely comprised of the peripheral vomeronasal organ (VNO), the accessory olfactory bulb (AOB) and the medial subnucleus of the amygdala (MeA), we studied the molecular diversity and neuronal subtype connectivity of this interconnected circuit. We show that populations of neurons of the AOS can be molecularly subdivided based on their current or prior expression of the transcription factors *Foxp2* or *Dbx1*. We show that the majority of AOB neurons that project directly to the MeA are of the *Foxp2*-lineage. Using single cell patch clamp electrophysiology, we further reveal that in addition to sex-specific differences across lineage, the relative contributions of excitatory and inhibitory inputs to MeA *Foxp2*-lineage neurons differ between sexes. Together, this work uncovers a novel molecular diversity of AOS neurons and lineage- and sex-differences in patterns of connectivity.

## 1. Introduction

### 1.1 Rodent Olfactory Circuitry

The olfactory system is subdivided into two functionally distinct components: the main olfactory system (MOS) and the accessory olfactory system (AOS). Odorants that activate the MOS bind to receptors in the main olfactory epithelium (MOE) in the nose, which projects directly to the main olfactory bulb (MOB) in the brain (Dulac & Wagner, 2006). From here information is sent to higher order brain regions, primarily the olfactory and piriform cortices (Hintiryan et al., 2012; Shipley & Adamek, 1984). The AOS processes both volatile and non-volatile cues that are mainly dedicated for innate behaviors such as mating, territorial defense and predator avoidance (Papes et al., 2010). Non-volatile cues include pheromones, for example, those released by anal and lacrimal glands and those present in urine (Cavaliere et al., 2020; Stowers & Liberles, 2016). These cues impart information regarding the hormonal state and sex of a conspecific (Dulac & Wagner, 2006; Thoß et al., 2019). Odorants that stimulate the AOS bind directly to receptors in the vomeronasal organ (VNO), which is located in the lower part of the nasal septum, in proximity to the roof of the mouth. VNO olfactory sensory neurons project directly to the accessory olfactory bulb (AOB), which anatomically sits apart from the MOB in the posterior-dorsal aspect of the olfactory bulb (OB) (Wagner et al., 2006). The AOB projects directly to the medial (MeA) and cortical (CoA) nuclei of the amygdala and the bed nucleus of stria terminalis (BNST), all of which are interconnected and send robust projections to the hypothalamus (Dulac & Wagner, 2006). The direct input from the MOB and AOB to higher-order processing centers in the brain is unique to the olfactory system as all other sensory modalities (touch, taste, vision, hearing) are first relayed through the thalamus. Thus, the main and accessory olfactory systems work in parallel and in tandem to allow an animal to interpret its complex olfactory world rapidly without thalamic processing.

### 1.2 Olfactory System Neuronal Diversity

The diversity of olfactory cues that an animal senses in its environment is vast and complex. This is reflected in the large number of receptors; hundreds and thousands, that are present in the sensory neurons of the VNO and MOE, respectively. The logic by which this complex sensory information is processed in the brain is currently much better understood in the MOS than the AOS. In the MOS, there appears to be regionalization of olfactory cue identification in the MOE and subsequent sorting in the MOB. The MOE is subdivided into multiple anatomically and molecularly distinct zones which recognize different odorant classes (Ruiz Tejada Segura et al., 2022). This segregation is maintained in the MOB, which is also subdivided based on the nature of the cue (Burton et al., 2022; Sakano, 2010). Based on cell morphology, anatomic sublocalization and physiological properties (Nagayama et al., 2014), the mitral and tufted (M/T) output neurons of the MOB are heterogeneous. More recent RNA-seq studies have uncovered a deeper level of diversity of MOB output neurons, identifying at least 8 distinct molecular subtypes (Zeppilli et al., 2021). This molecular coding in the MOB appears to predict neuronal subtype-specific patterns of inputs to higher-order neurons in the piriform cortex, although this remains an area of intense investigation (Adam et al., 2014; Uchida et al., 2014).

Although less well-characterized than the MOS, neurons within the VNO and AOB also appear to be diverse by several criteria. The VNO is subdivided into anatomically and molecularly distinct apical and basal layers, which differentially project to the anterior (aAOB), and the posterior (pAOB) AOB, respectively (Knöll et al., 2003). Interestingly, the aAOB and pAOB are also molecularly distinct from each other (Huilgol et al., 2013), and may regulate different types of innate behaviors (e.g. reproductive versus aggressive) (Kumar et al., 1999; Montani et al., 2013; Nunez-Parra et al., 2011). Like in the MOB, M/T cells comprise the output neuron populations of the AOB. Based on morphological criteria, there appear to be at least 3 subtypes of AOB output neurons (Larriva-Sahd, 2008; Yonekura & Yokoi, 2008). However, the molecular diversity of both VNO and AOB output neurons remains unexplored.

In the MeA, the direct synaptic target of AOB output neurons, prior work from our lab and others has revealed up to 20 different neuronal subtypes, as characterized by their molecular and/or intrinsic electrophysiological properties (Bian, 2013; Carney et al., 2010; Chen et al., 2019; Keshavarzi et al., 2014; Lischinsky et al., 2017; Matos et al., 2020). In addition to possessing a variety of local interneuronal subtypes, the MeA is comprised of both excitatory and inhibitory output neurons (Bian et al., 2008; Choi et al., 2005; Wu et al., 2017). These MeA output neurons can be further subdivided based on their diverse morphological, electrophysiological and/or molecular properties (Keshavarzi et al., 2014; Lischinsky et al., 2017; Matos et al., 2020), which additionally appears to predict behavioral regulation (Lischinsky et al., 2022). Molecular diversity within the MeA is defined by differential expression of transcription factors (e.g. *Otp*, *Foxp2* and *Dbx1*) in progenitors that give rise to different output classes. These classes display different patterns of expression of select neurohormones (e.g. Aromatase, estrogen and androgen receptors) (Lischinsky et al., 2017) and ion channels (e.gs. *Kv7.1, Kir5.1, Kir2.1, Slo2.2*) (Matos et al., 2020). This parcellation of MeA identity by molecular expression raises the question of whether other components of the interconnected AOS, such as the VNO and AOB, may be subdivided by the expression of similar molecular identifiers; a question which we address in experiments described herein.

### 1.3 Sex differences in the AOS

One of the major functions of the AOS is to process information as to the reproductive state of conspecifics (Ben-Shaul et al., 2010). This information appears to be encoded differently in male and female brains. For example, cues from an estrus female elicit behavioral responses that are very different depending on whether the cue is being detected by a female or a male mouse. Although much remains to be understood, the first level of distinguishing between same or opposite sex cues is likely initiated in the VNO, where there are dedicated receptors for sex-specific cues (Isogai et al., 2011; Li & Dulac, 2018). Beyond this peripheral parsing of male/female cues, the processing of sex-specific information also occurs higher-order brain regions, such as the MeA and BNST (Bergan et al., 2014; Li et al., 2017; Rigney et al., 2019). However, how this information is relayed and processed in the MeA remains an open question.

The MeA displays extensive sex differences in the intrinsic properties of cell morphology, dendritic complexity, cell size, intrinsic biophysical properties, and gene expression (Cooke et al., 2007; Cooke & Woolley, 2005; Hines et al., 1992; Matos et al., 2020). Using cFos staining (Lischinsky et al., 2017) and *in vivo* neuronal population recordings, our work and the work of others revealed that the MeA also displays sex differences in neuronal population responses to olfactory cues. Patch clamp single neuron electrophysiological and neuronal tracing studies have also revealed sex differences in patterns of inputs to the MeA. For example, neurons in the male MeA display more excitatory input than females (Billing et al., 2020; Cooke & Woolley, 2005). However, the MeA neuronal subtype target of these inputs remains unknown. Here, in addition to studying lineage diversity in the AOS, we took advantage of our ability to specifically tag two major populations of MeA GABAergic output neurons, *Foxp2*- and *Dbx1*-lineage cells, to further address putative sex differences in synaptic inputs. Such information is necessary to piece together a neuronal subtype-level understanding of how male and female brains differentially process olfactory information for appropriate behavioral outputs.

### 1.4 Summary

The identification of neuronal diversity by early transcription factor expression has been essential to understanding the functionality of neurons across the nervous system. In particular, this approach has been highly informative in unraveling the function of cortical interneurons (Batista-Brito et al., 2008; Mayer et al., 2018; Mi et al., 2018). Here, we used transcription factor expression, specifically current or prior expression of *Foxp2* or *Dbx1* as an entry-point to parse neuronal diversity in the AOS, and further explore sex-specific patterns of input to the MeA, a main locus of convergence for olfactory cues with innate behavioral relevance. Our findings also provide a platform for future exploration into the specific behavioral roles played by transcription-factor identified AOS circuits.

## 2. Materials and Methods

### 2.1 Animals

Mice were housed in a 12h light/dark cycle and had *ad libitum* access to food and water. All mice used here were considered adult at > 2 months, with specific ages for each experiment indicated below. Mice used were: *Foxp2^cre^* (Jax 030541), *Dbx1^cre^* (Bielle et al., 2005), *C57BL/6* (JAX #000664), *LSL-FlpO* (JAX #028584), and *Rosa26YFP* (JAX# 006148). Mice were genotyped using a commercial genotyping service (Transnetyx Inc., Cordova, TN). *Foxp2^cre^* and *Dbx1^cre^* mice were maintained as heterozygotes on the C57BL/6 background and crossed to *Rosa26YFP* mice as described in the Results Section.

### 2.2 Viruses and Stereotaxic Surgery

All experimental and surgical procedures were conducted in accordance with and approved by the Institutional Animal Care and Use Committee at Children’s National Hospital. For all surgeries, adult (2-5 month old) animals were anaesthetized with isoflurane and placed into a stereotaxic apparatus (Stoelting Co. #51600). Body temperature was maintained with a heating pad during surgery and recovery, and 1.5–2% isoflurane was delivered continuously through a nose port. Animals were treated with analgesic buprenorphine (0.09 mg/kg body weight, of 0.03 mg/mL buprenorphine prepared in sterile saline) prior to surgery, and every 12 hours afterwards, as needed. Mice were monitored daily and sacrificed 2-5 weeks post viral infection.

#### Accessory Olfactory Bulb

400 nL at 30 nL/min. of *AAV5-hSyn-Con/Foff.EYFP.WPRE* and *AAV5-hSyn-Coff/Fon.EYFP.WPRE* at a titer of 2.6 X 10^12^ GC/mL (University of North Carolina Vector Core), combined 1:1 with blue fluorescent 1% solid polymer microspheres (Thermo Fisher Scientific, Cat#B0100) were injected unilaterally into the accessory olfactory bulb (AOB) (AP: +4.1; ML: ±1.0, DV: -1.5 from Bregma) with a Hamilton syringe into either *Foxp2^cre^* (*AAV5-hSyn-Con/Foff.EYFP.WPRE)* or *Dbx1^cre^;FlpO* (*AAV5-hSyn-Coff/Fon.EYFP.WPRE*) adult mice. Following virus delivery, the syringe was left in place for 3 min to prevent backflow and then slowly withdrawn.

#### Medial Amygdala

200 nL at 15 nL/min. of AAV-retro (*retroAAV2-Ef1a-DO_DIO-TdTomato_EGFP-WPRE-pA*) were injected bilaterally into the MeA (AP: -1.6, ML: ±2.2, DV: -4.8 from Bregma) of adult mice.

### 2.3 Immunohistochemistry

Animals were anaesthetized and transcardially perfused with 10 mL of 1X Phosphate Buffered Saline (PBS) followed by 10 mL of 4% paraformaldehyde (PFA) in 1X PBS. After perfusion, the brains were extracted and incubated overnight in the same fixative and cryoprotected in phosphate-buffered 30% sucrose solution for >48h at 4^0^C. After cryoprotection, brains were embedded in O.C.T Compound (Fisher HealthCare Cat No. 23-730-571). Serial 40-60 μm coronal cryosections were cut using a cryostat (CM3050S, Leica) and collected in PBS containing 0.02% sodium azide. Sections were then incubated in blocking buffer (10% normal donkey serum (NDS), 0.5% Triton X-100, 1X PBS) for 1h at room temperature (RT) and subsequently incubated in the primary antibody mixture diluted in blocking buffer overnight at 4 °C. The primary antibodies were: Rabbit anti-Foxp2 (1:1000, Atlas Antibodies, Cat No. HPA000382), Rat anti-GFP (1:1000, Nalacai Tesque Inc, Cat No. 04404-84), Rabbit anti-CART (1:500, Phoenix Pharmaceutical Inc, Cat No. H003-62), Rabbit anti-PDE4 (Abcam Cat No. ab14607), goat anti-OMP (Wako, Cat No. 544-10001), Rabbit anti-PSD95 (1:50, Proteintech, Cat No.20665.AP), Guinea Piganti-VGLUT2 (1:100, EMD Millipore, Cat No. AB5905), Mouse anti-VGAT (1:1000,Invitrogen, Cat MA5-2463), Rabbit anti-Gephyrin (1:200, Thermofisher, Cat PA5-5517). Sections were rinsed 3 × 10 min with PBST (1X PBS with 0.5% Triton X-100) and incubated in appropriate secondary antibodies (Jackson ImmunoResearch) conjugated to Alexa 488, Alexa 594, or Alexa 647 fluorescent dyes for 1h at RT. Sections were washed 3 × 10 min with PBST and 1 x 10 min with 1X PBS. Slides were mounted with Fluoromount G® with DAPI (Thermo-fisher, Cat No.0100-20).

### 2.4 Multiplexed fluorescent *in situ* hybridization

8-week old male *Dbx1^cre^;RYFP* mice were transcardially perfused with 10 mL ice-cold 1X PBS, followed by 10 mL freshly-prepared ice-cold 4% paraformaldehyde in 1X PBS. Whole brains were collected, incubated in 4% PFA/1X PBS at 4^0^C for 24h, followed by cryoprotection in 30% sucrose in 1X PBS for 48h at 4^0^C. Brains were then embedded in O.C.T. compound and stored at -80^0^C until cryosectioning. 10 µm slide-mounted cryosections of the medial amygdala from Bregma -1.58 mm to Bregma -1.94 mm (identified by comparison to the mouse brain atlas, Franklin & Paxinos, 2008) were obtained using a Thermo Scientific HM525 NX cryostat at – 22^0^C. Slides were stored at -80^0^C with dessicant until staining. RNAscope^TM^ HiPlex-12 Mouse kit (ACDBio, Cat #324106) was used to detect the following *Mus musculus* mRNA targets, available through ACDBio’s catalog: *Tac2* (#446391), *Tacr1* (#428781), *Trh* (#436811), *Foxp2* (#428791), *Avp* (#401391), *Ucn3* (#464861), *Npy1r* (#427021), *Htr2c* (#401001), *Npy* (#313321), *Ecel1* (#475331) and *EYFP* (#312131). Protocol for HiPlex-12 sample preparation, pretreatment and staining was followed exactly as per ACDBio’s user manual for fixed, frozen tissue sections.

### 2.5 Microscopy

To analyze the labeled cells in the AOB and output projections to the MeA, 10X and 20X epifluorescence images were acquired using Olympus BX63. All off-target regions were excluded from analysis. All HiPlex-12 samples were imaged at a Leica DMi8 THUNDER deconvolution microscope using 10X (NA 0.32) and 63X oil (NA 1.40) objectives. 395, 470, 550 and 640 nm excitation lines and corresponding emission filter sets (440, 510, 590 and 700 nm respectively) were used. 20-200 ms camera exposure time was used for each channel, varied depending on observed signal-to-noise in that channel and maintained uniformly across samples. THUNDER large-volume adaptive deconvolution with computational clearing was performed on acquired widefield images. Deconvolution settings: 1.47 refractive index, Good’s Roughness method of regularization with parameter 0.05, medium optimization, number of iterations and cutoff gray value set to ‘Auto’, feature scale of 535 nm (minimum possible) for all channels except DAPI, feature length of 2673 nm (software default) for DAPI, 98% strength for all channels. Single-Z plane tile-scan images of the MeA were acquired with automatic linked shading correction, deconvolved, stitched (statistical blend) and exported as TIFFs for further image processing and quantification.

### 2.6 Patch clamp electrophysiology recordings

Sexually naive, adult mice (P50–P90) were deeply anaesthetized with isoflurane and sacrificed. Brains were removed and immediately immersed in an ice-cold carbogenated (95% O_2_ and 5% CO_2_) sectioning solution (75 mM sucrose, 10 mM D-glucose, 25 mM NaHCO_3_, 87 mM NaCl, 2.5 mM KCl, 1.0 mM NaH_2_PO_4_, 1.0 mM MgCl_2_ hexahydrate, and 0.5 mM CaCl_2_ dihydrate; pH 7.3; 295-300 mOsm/kg. 300 µm coronal slices were sectioned on a vibratome (Leica VT1200S) at the level of posterior MeA (Bregma -1.56 to -1.94 mm; Franklin and Paxinos, 1997). Slices were collected and placed in oxygen-equilibrated artificial CSF (ACSF) composed of the following: 125.0 mM NaCl, 3.5 mM KCl, 1.0 mM MgCl_2_ hexahydrate, 1.25 mM NaH_2_PO_4_, 2.0 mM CaCl_2_ dihydrate, 26.0 mM NaHCO_3_, and 10.0 mM D-glucose; pH 7.3; 295–300 mOsm/kg. *Dbx1^cre^;RYFP*-positive or *Foxp2^cre^;RYFP*-positive neurons were visualized using a Nikon FN1 epifluorescence microscope with a 450 to 490 nm filter. Whole-cell patch-clamp recordings from YFP-positive fluorescent cells were performed at RT with continuous perfusion of carbogenated ACSF. Signals were acquired on a patch-clamp amplifier (Multiclamp 200B) and digitized at 250 kHz with an A/D converter (DigiDATA1550B). Recordings were performed with glass electrodes pulled on a Sutter P-2000 pipette puller (Sutter Instruments), with 3.5 MΩ resistance and filled with a potassium gluconate-based intracellular solution containing the following: 119.0 mM K-gluconate, 2.0 mM Na-gluconate, 6.0 mM NaCl, 2.0 mM MgCl_2_ hexahydrate, 10.0 mM HEPES, 0.9 mM EGTA, 4.0 mM Mg-ATP, 14.0 mM Tris-creatine PO_4_, and 0.3 mM Tris-GTP; pH 7.3; 285–295 mOsm/kg. Cells were first recorded in current clamp configuration as reported in Matos et al. (2020). Subsequently, they were switched from current clamp to voltage clamp configuration. Each seal was tested to have maintained access resistance at < 30 MΩ throughout the current and voltage clamp recordings, verified at the beginning and end of a recording session. Cells that did not meet these criteria were excluded from all experiments. Voltage clamp recordings were done in three progressive stages, for 3 minutes at each stage: voltage clamp gap free holding at -60 mV in ACSF only; voltage clamp gap free holding at -60 mV in ACSF + 50 µM Picrotoxin (to block GABAARs); and voltage clamp gap free holding at -60 mV in ACSF + 50 µM Picrotoxin + APV (to block GABAARs and NMDARs). Only the 3^rd^ minute of recording for each stage was analyzed to ensure that each successive drug had exerted its full effect. All measurements were acquired and analyzed offline using Clampfit Software 10.6 (Molecular Devices). Statistical analysis and plotting of event frequency data were performed in GraphPad Prism 9. Data were determined to have a lognormal distribution using D’Agostino-Pearson’s test, and then log transformed for subsequent analyses with 2-way ANOVA using sex and lineage as independent variables. Homoscedasticity and normality of residuals of log transformed data were verified by Spearman’s test and D’Agostino-Pearson omnibus (K2) tests respectively. ANOVA interaction and main effects were inspected (α=0.05) using F test, followed by Sidak’s test for multiple pairwise planned comparisons with α=0.05.

### 2.7 Image Processing, Quantification and Statistical Analyses

All immunohistochemical images analysis was done using Fiji (Schindelin et al., 2012). For *in situ* hybridization data, linear brightness/contrast adjustments were made for all channels in Fiji similarly across tissue sections, ROIs for the MeA were drawn based on Franklin & Paxinos, (2008) mouse brain atlas (using 10X tile-scanned DAPI image of whole section) and markers were added to Foxp2+ and EYFP+ cells (DAPI+ nuclei with at least 6 mRNA puncta on or clustered tightly around nucleus) in the MeA using the Cell Counter plugin. These were aligned individually to images of each investigated mRNA target using DAPI channel for transparency-based manual image registration in Adobe Photoshop. MeA cells expressing each marker (6 or more mRNA puncta on or tightly clustered around nucleus) were quantified within *Foxp2*+ and *EYFP*+ cells and statistically analyzed (Mann-Whitney U tests followed by FDR correction for multiple comparisons using Benjamini-Hochberg procedure with 10% FDR) and plotted using GraphPad Prism 9.

## 3. Results

### 3.1 Foxp2+ and *Dbx1*-lineage cells in the VNO and AOB

Our previous studies revealed that Foxp2+ and *Dbx1*-lineage neurons comprise large, non-overlapping populations of neurons in the MeA (Lischinsky et al., 2017). In the MeA, Foxp2 is first expressed during embryogenesis and remains on through adulthood. In contrast, *Dbx1* is expressed only in mitotic forebrain ventricular zone (VZ) progenitors during embryogenesis and is turned off when they transition to the subventricular zone (SVZ) (Hirata et al., 2009). To mark *Dbx1*-lineage neurons we used an anti-GFP antibody on tissue from previously validated *Dbx1^cre^*mice crossed to *RYFP* reporter mice **(Figures 1, 2, 5, 6 & 7)**. For gene expression studies (**Figures 2, 5 & 6**), where we assessed both *Dbx1*-lineages and Foxp2+ cells, Foxp2+ neurons were identified using a well-characterized antibody or by RNA scope *in situ* hybridization. For viral Cre-based connectivity tracing, electrophysiology and synaptic experiments (**Figures 3, 4, 7, 8 & 9)**, we used previously validated *Foxp2^cre^* mice (Rousso et al., 2016) crossed to *RYFP* reporter mice or injected with an *EYFP*- and/or *TdTomato*-carrying reporter virus.

**Figure 1:**
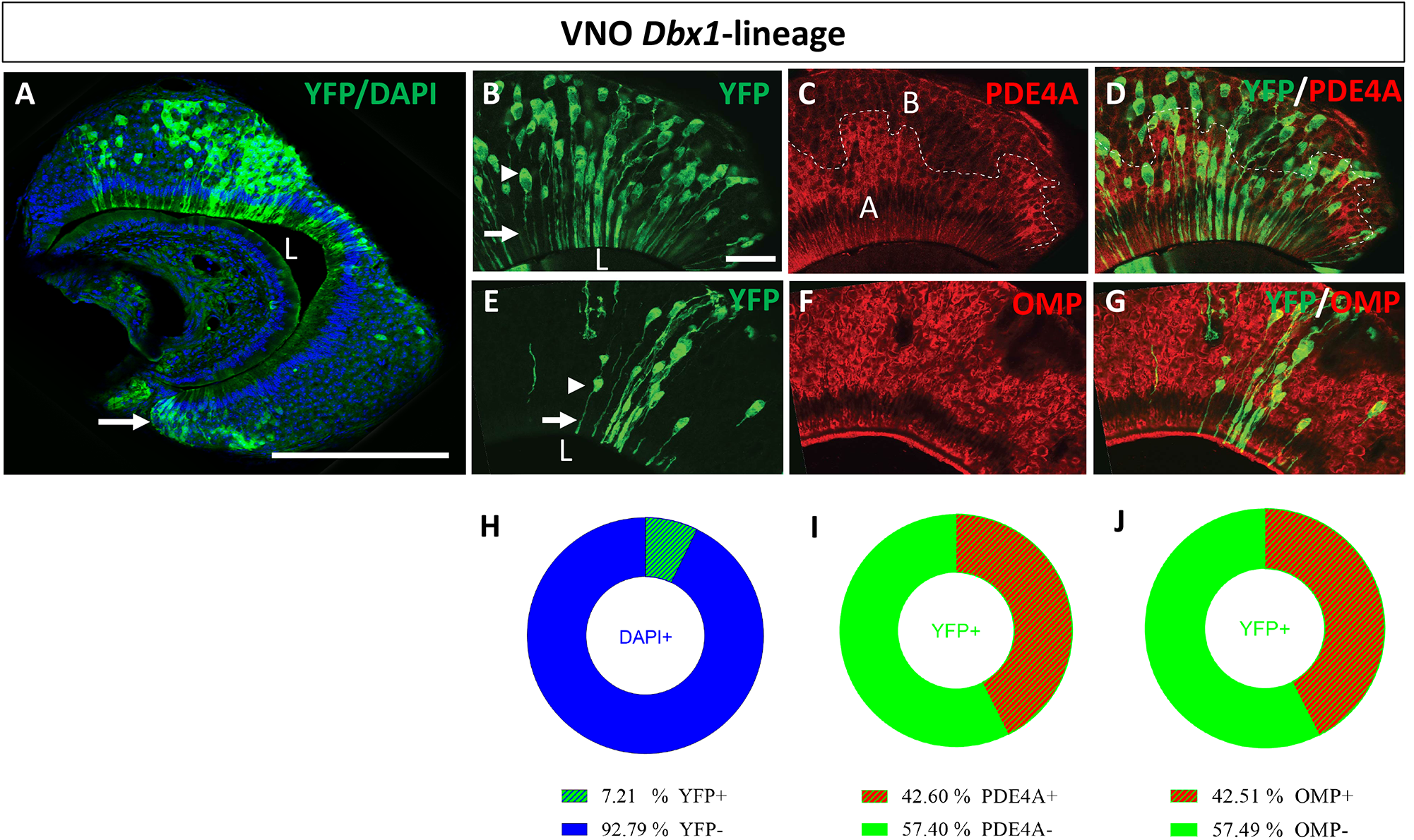
*Dbx1*-derived chemosensory neurons in the VNO. (**A**) Coronal section through the VNO of a *Dbx1^cre^;RYFP* mouse shows recombined YFP+ neurons (green) as a subset of the total DAPI+ (blue) cell population. Arrowhead marks a cluster of *Dbx1*-lineage cells in the putative neurogenic niche. (**B**) Co-immunostaining for YFP (green) and (**C**) PDE4A (red), which marks the apical (A) layer of the VNO. Arrowhead and arrow in (**B**) denote cell body and dendritic process respectively, projecting to the lumen (L) of a *Dbx1*-lineage olfactory sensory neuron. (**D**) Merged image of YFP and PDE4A reveals *Dbx1*-lineage cell bodies in both the apical and basal layers. (**E, F**) Co-immunostaining for YFP (green, **E**) and OMP (red, **F**), which marks mature sensory neurons. (**G**) Merged image of YFP and OMP reveals mature *Dbx1*-lineage neurons. (**H-J**) Donut charts show quantification of percent contribution of *Dbx1*-lineage cells to the whole VNO population (**H**), to the PDE4A+ apical domain and the PDE4A-basal domain (**I**), and the OMP+ mature and OMP-immature populations (**J**). Scale bars: 200 µm (**A**), 20 µm in (**B-G**). n=18 mice (5-18 sections/mouse) in (**H-J**).

**Figure 2:**
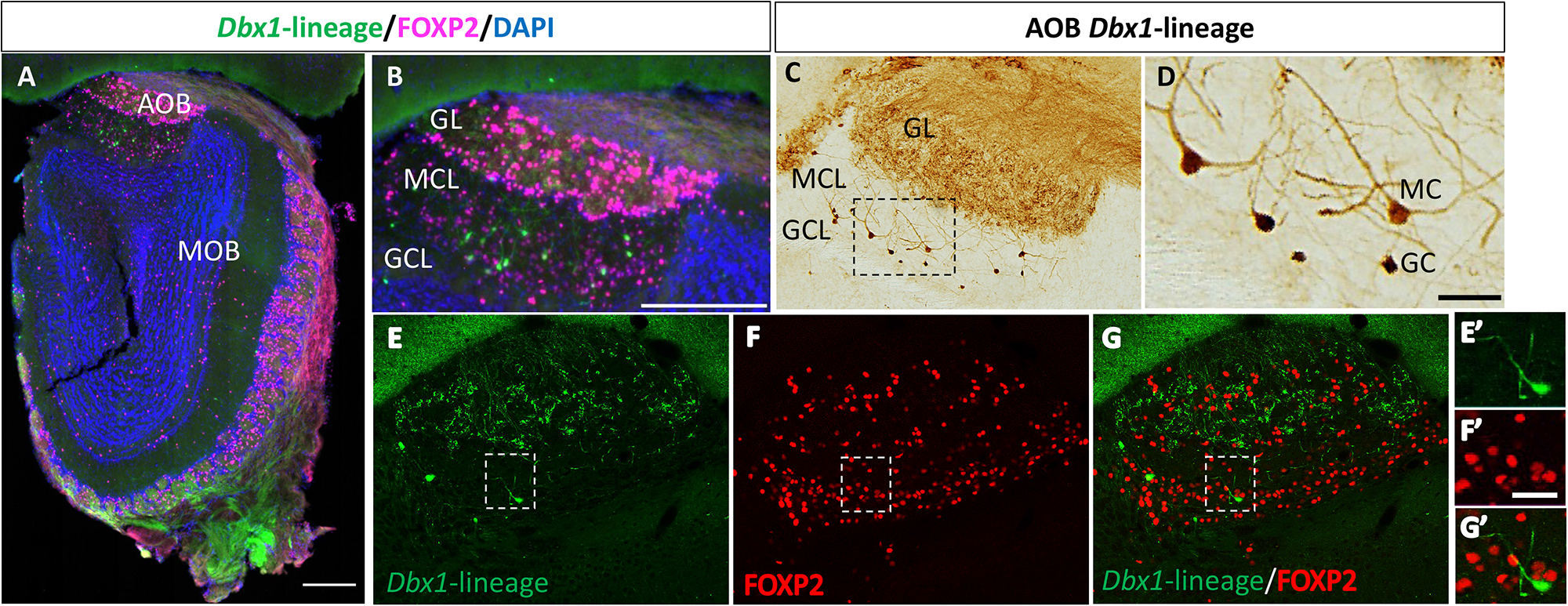
*Dbx1*-derived and Foxp2+ neurons in the AOB. (**A**) Low power view of a coronal section from the olfactory bulb (OB) reveals many Foxp2 immunopositive cells (magenta) throughout both the accessory olfactory bulb (AOB) and main olfactory bulb (MOB), with fewer *Dbx1*-lineage immune-positive cells (green). (**B**) Higher magnification view shows Foxp2+ cells through all layers of the AOB, with *Dbx1*-lineage cells present mainly in the mitral cell layer (MCL) and granule cell layer (GCL). (**C**) Many YFP+ fibers are observed in the GL, likely from *Dbx1*-lineage sensory neurons in the VNO. Boxed region highlights a handful of *Dbx1*-lineage neurons in the MCL and GCL. (**D**) Higher magnification of the boxed region reveals morphologies of *Dbx1*-lineage putative MCs and GCs. (**E-G**) In the AOB, *Dbx1*-lineage (green) and Foxp2+ (red) are non-overlapping populations. (**E’-G’**) High power magnification highlighting a single *Dbx1*-lineage cell within a population of Foxp2+ cells. Scale bar in A equals 250 µm, Scale bar in **B** (also for **E, F, G**) equals 250 µm, Scale bar in **D** equals 20 µm, scale bar in **F’** equals 20 µm (also for **E’** and **G’**).

**Figure 3:**
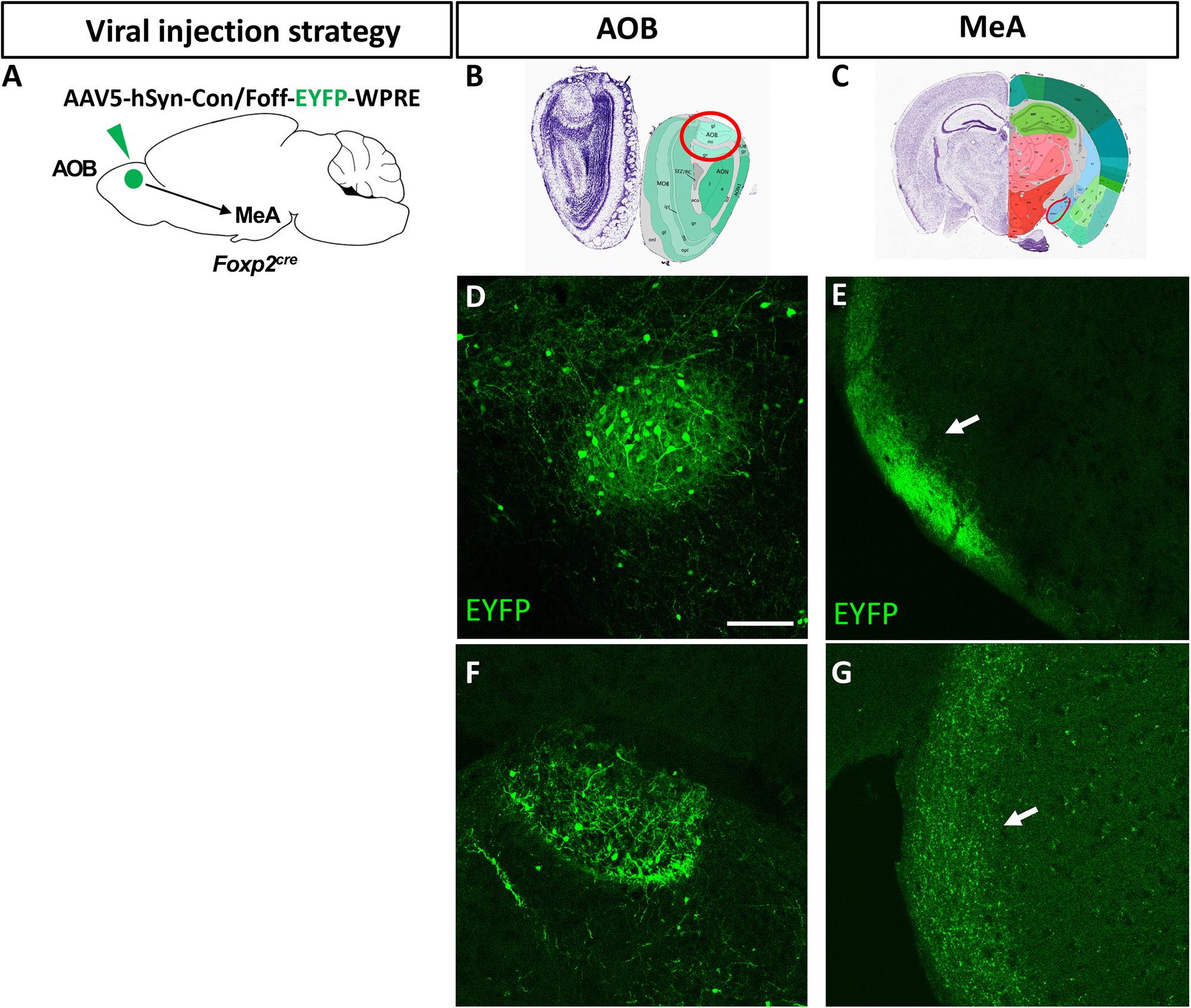
AOB Foxp2+ cells project to the MeA. (**A**) Schematic of the strategy of anterograde viral infection into the AOB of *Foxp2^cre^* mice. (**B, C**) Images from the Allen Mouse Brain Atlas: https://help.brain-map.org/display/mousebrain/Documentation highlighting the AOB and MeA, shown in (**D, F**) and (**E, G**), respectively. (**D, F**) Representative coronal sections of virally infected AOB (n=17 successful infections) showing recombined EYFP+ neurons (green) in the AOB. (**E, G**) EYFP+ axons (arrows) from AOB infected cells observed along the input path from the AOB to the MeA. Scale bar, 100 µm.

**Figure 4:**
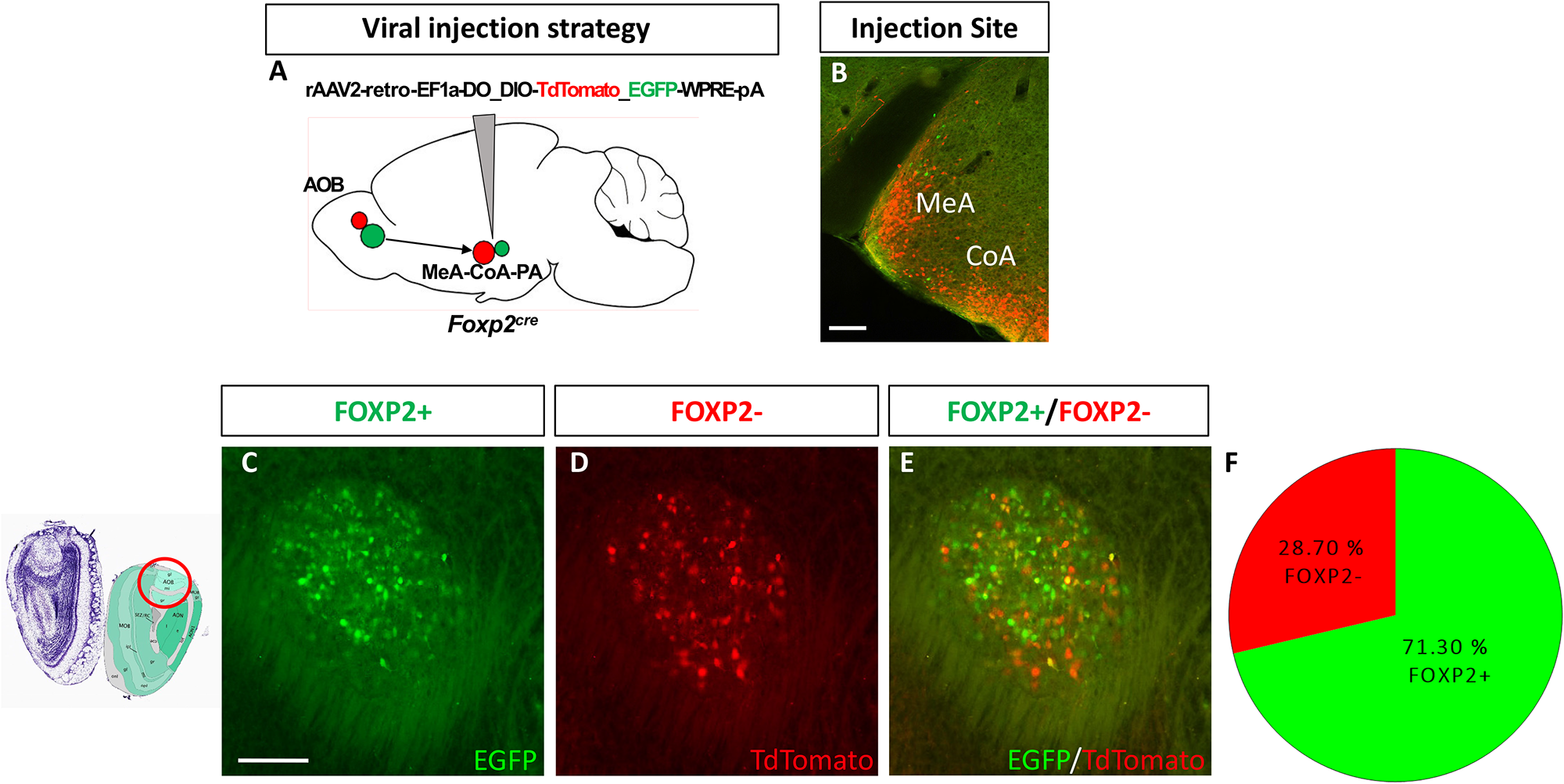
AOB inputs to MeA *Foxp2*-derived neurons. (**A**) Schematic illustrating strategy of retrograde viral injection into the MeA-CoA-PA with a dual retrograde AAV tracer in a *Foxp2^cre^* mouse. (**B**) Immunohistochemistry (coronal view), at the injection site (MeA-CoA) allows visualization of targeted region. Image from Allen Mouse Brain Atlas: https://help.brain-map.org/display/mousebrain/Documentation shows region of analysisin coronal sections through AOB. (**C-E**) Immunohistochemistry shows recombination of intermingled Foxp2+ (EGFP+, green) (**C**) and Foxp2-(tdTomato+, red) (**D**) cells in coronal views of the AOB (merge, **E**). (**F**) Pie chart showing quantification of the percentage of Foxp2+ (green) and Foxp2-(red) cells sending input from the AOB to MeA-CoA-PA. Scale bar: 200 µm in (**B-E**). n=7 mice (10-17 sections/mouse).

**Figure 5:**
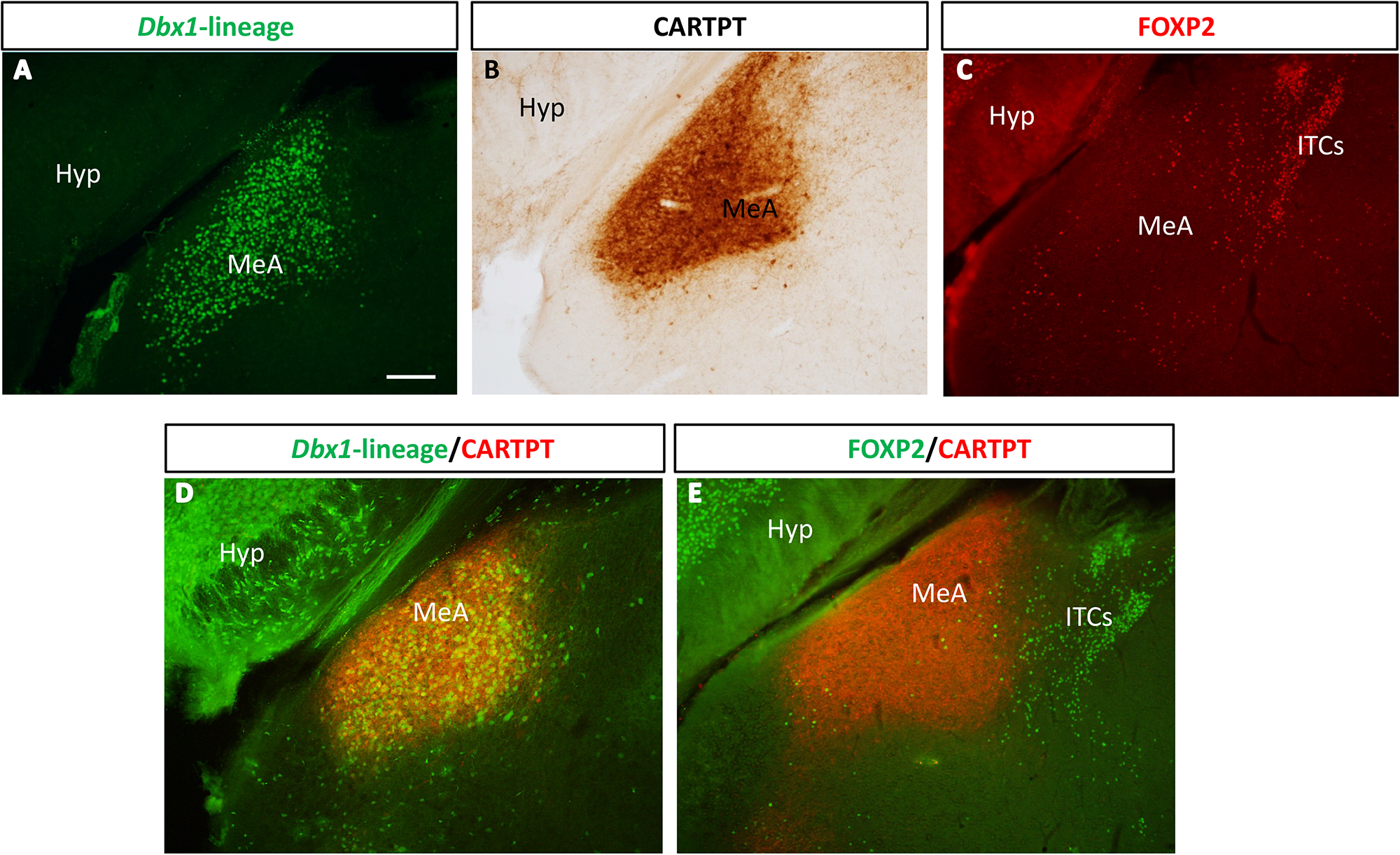
Distribution of *Dbx1*-lineage and Foxp2+ neurons in the MeA in relation to CARTPT expression. (**A**) *Dbx1*-lineage neurons (green) are present in a tight cluster in the posterior MeA as shown in coronal sections. (**B**) Expression of CARTPT, which marks both cell bodies and incoming axonal terminals, matches the distribution of *Dbx1*-lineage neurons. (**C**) The distribution of Foxp2+ neurons (red) is complementary to the expression pattern of CARTPT. (**D**) *Dbx1*-lineage neurons (green) are embedded within a zone of CARTPT (red) expression. (**E**) Foxp2+ cells (green) are observed predominantly outside the zone of CARTPT expression (red). Abbreviations: Hyp, hypothalamus; MeA, Medial Amygdala; ITCs, amygdala intercalated clusters. Scale bar, 200 µm.

We first determined if Foxp2+ or *Dbx1*-lineage cells are present in the VNO (**Figure 1**) and AOB (**Figure 2**) of postnatal (1-3 months of age) mice. The adult VNO is comprised of mature olfactory receptor neurons (ORNs) and immature newly generated ORNs in a neurogenic niche located at the edge of the VNO (Katreddi & Forni, 2021). While no Foxp2+ cells were found in the adult VNO (data not shown), we observed numerous *Dbx1*-lineage cells, comprising ∼7% of the total DAPI+ population across the entire VNO (**Figure 1A, H**). The adult VNO is comprised of a lumen-facing apical layer and a non-lumen-facing basal layer. These layers project to the anterior and posterior AOB, respectively (Knöll et al., 2003). The apical layer is marked by PDE4A, a member of the cAMP-specific family of phosphodiesterases (Lau & Cherry, 2000). We found *Dbx1*-lineage ORN cell bodies almost evenly distributed across the PDE4A+ (∼43%) apical layer and the PDE4A-(∼57%) basal layer (**Figure 1B-D**, **I**). Co-labeling of YFP+ cells with OMP, a marker for mature VNO neurons (Farbman & Margolis, 1980) revealed that ∼43% of *Dbx1*-lineage neurons are mature (**Figure 1E-G, J**). Thus, in the VNO, *Dbx1*-lineage cells comprise both immature and mature ORNs located in both the apical and basal layers.

In the olfactory bulb (OB), we next examined if Foxp2+ and *Dbx1*-lineage cells were present in the MOB and the AOB. Co-immunostaining for Foxp2 and YFP in *Dbx1^cre^;RYFP* mice revealed cells from both populations across the OB (**Figure 2A**), with an apparent greater number of Foxp2+ cells. Focusing on the AOB, which receives direct input from the VNO, we observed Foxp2+ and *Dbx1*-lineage cells located within the glomerular, mitral and granular cell layers (**Figure 2B-G**). The expression of YFP in axons also allowed us to observe putative projections from the *Dbx1*-lineage VNO ORNs. This was best visualized by DAB immunostaining in *Dbx1^cre^;RYFP* mice. where observed a strong bundle of YFP+ fibers projecting into the AOB (**Figure 2C**). Across all layers of the AOB, we found that Foxp2+ cells represented a larger population than *Dbx1*-lineage cells. We further found that in the mitral and granule cell layers which comprise the AOB output neurons, the majority of Foxp2+ and *Dbx1*-lineage cells were separate populations (only 1%, 53/5217 cells co-express Foxp2 and YFP; **Figure 2G, G’**), a population segregation that mimics what we previously observed in the MeA (Lischinsky et al., 2017).

### 3.2 *Foxp2*-lineage neurons comprise the majority of MeA-projecting AOB output neurons

The MeA receives direct projections from the AOB (Zheng et al., 2020). We next wanted to examine whether *Foxp2*- and *Dbx1*-lineage neurons in the AOB project to the MeA. To accomplish this, we injected an anterograde *AAV5-hSyn-Con/Foff.EYFP.WPRE* virus into the AOB of *Foxp2^cre^* mice (**Figure 3A**) and an anterograde *AAV5-hSyn-Coff/Fon.EYFP.WPRE* virus into *Dbx1^cre^;FlpO* mice. Infections in *Dbx1^cre^;FlpO* mice resulted in recombination of only a few *Dbx1*-lineage neurons, therefore we were not able to trace their projections (data not shown). This low number of recombined AOB cells is likely a reflection of the low number of *Dbx1*-lineage M/T cells as compared to Foxp2+ cells (**Figure 2**). In contrast, in infections of *Foxp2^cre^* AOB cells, we observed large numbers of recombined cells (**Figure 3B, D, F**), with robust projections emanating from the AOB to the MeA (**Figure 3C, E, G**). Outputs were assessed by co-labeling with GFP to identify axon tracts.

As we observed strong projections from *Foxp2*-lineage AOB M/T neurons to the MeA, we next wanted to determine if this population represented most of the projections to the MeA. To accomplish this, we injected a dual reporter retrograde ‘retro-AAV2’ virus (*retroAAV2-Ef1a-DO_DIO-TdTomato_EGFP-WPRE-pA*) (Tervo et al., 2016) into the amygdala of *Foxp2^cre^* mice (**Figure 4A, B**). Retro-AAV2 is taken up by presynaptic neuronal terminals and translocated retrogradely to cell bodies where presynaptic Cre-expressing cell bodies are EGFP+/TdTomato-while Cre-negative cell bodies are EGFP-/TdTomato+. In the AOB, we found both EGFP+ and TdTomato+ cells (**Figure 4C-E**) indicating that both Foxp2+ and Foxp2-AOB M/T neurons project to the MeA. Quantification of the numbers of EGFP+ and TdTomato+ cells revealed that the EGFP+ population represented a majority of recombined neurons (**Figure 4F**). Thus, in the AOB, most of the amygdala-projecting neurons are of the *Foxp2*-lineage.

### 3.3 MeA Foxp2+ and *Dbx1*-lineage cells express different cohorts of neuropeptides

Our previous studies revealed that MeA Foxp2+ and *Dbx1*-lineage neurons express different neurohormones and ion channels (Lischinsky et al., 2017; Matos et al., 2020). Furthermore, a recent study has revealed that these lineages control different innate behaviors (Lischinsky, et al., 2022). The MeA expresses a variety of neuropeptides and receptors that likely play a neuromodulatory role in regulating innate behaviors such as mating, aggression, feeding, maternal care and social interaction, based on their known roles in other limbic nuclei. To explore whether these two populations express different combinations of neuropeptides we conducted immunohistochemistry (**Figure 5**) and multiplex RNAscope *in situ* hybridization (**Figure 6**) in sections from *Dbx1^cre^;RYFP* mice. Gene candidates were chosen based on the following criteria: The gene 1) is expressed in the MeA as shown either in prior published studies or the Allen Brain gene expression atlas (Lein et al., 2006), 2) plays a known role in MeA function or innate social behavior, and/or 3) was observed in a previous RNA-seq screen of the adult MeA (Chen et al., 2019). Following these criteria, we generated a list of 10 candidates (**Table 1**). The most striking pattern was the expression of CARTPT (Cocaine- And Amphetamine-Regulated Transcript Protein), a neuropeptide implicated in feeding, reward and stress (Carpenter et al., 2020; Funayama et al., 2022; Kristensen et al., 1998; Lee et al., 2022). We found CARTPT highly expressed in the MeA and in a pattern resembling the distribution of *Dbx1*-lineage cells (**Figure 5A, B**), and complementary to the distribution of Foxp2+ cells (**Figure 5C**). Dual immunofluorescence revealed that the majority of CARTPT+ cell bodies and projections were embedded within regions of *Dbx1*-lineage cells (**Figure 5D**). In contrast, CARTPT+ cell bodies and projections did not co-localize with Foxp2+ cells (**Figure 5E**). Thus, CARTPT expression appeared to be more associated with *Dbx1*-lineage cells than Foxp2+ cells. We next assessed the expression of the other 9 candidates by multiplexed RNAscope *in situ* hybridization in the MeA (**Figure 6**). Of these, we confirmed the expression of all candidates except *Npy1r*. Of the markers expressed in the MeA, most were expressed only in a small subset of either *Dbx1*-lineage or Foxp2+ cells. Of these markers, we found three viz. *Tac2, Ucn3* and *Npy* that were significantly expressed in more cells of one lineage versus the other. *Tac2* encodes a neuropeptide, tachykinin isoform 2, that promotes aggressive behavior in fruit flies and mice (Asahina et al., 2014; Zelikowsky et al., 2018).*Npy* encodes Neuropeptide Y which, in rodents, modulates aggression through Y1 receptors in the medial amygdala (Karl et al., 2004), is implicated in reduced social anxiety (Sajdyk et al., 1999, 2002) and regulates maternal behavior, a critical sex-specific social behavior (Muroi & Ishii, 2015). We found both *Tac2* and *Npy* mRNA were modestly but significantly enriched in *Foxp2+* cells relative to *Dbx1-*lineage cells in the MeA (**Figure 6A-A’’, C-C’’**). This is consistent with a role for *Foxp2*+ neurons in the MeA in intermale territorial aggression (Lischinsky et al., 2017, 2022). In contrast, we found *Ucn3* mRNA which encodes the Urocortin 3 peptide, enriched in *Dbx1*-derived cells in the MeA (**Figure 6B-B’’**). Urocortin 3 has been shown to promote preference for social novelty through its action in the MeA (Shemesh et al., 2016) and infant-directed aggression through its function in the perifornical area of the hypothalamus (Autry et al., 2021).

**Table 1:**
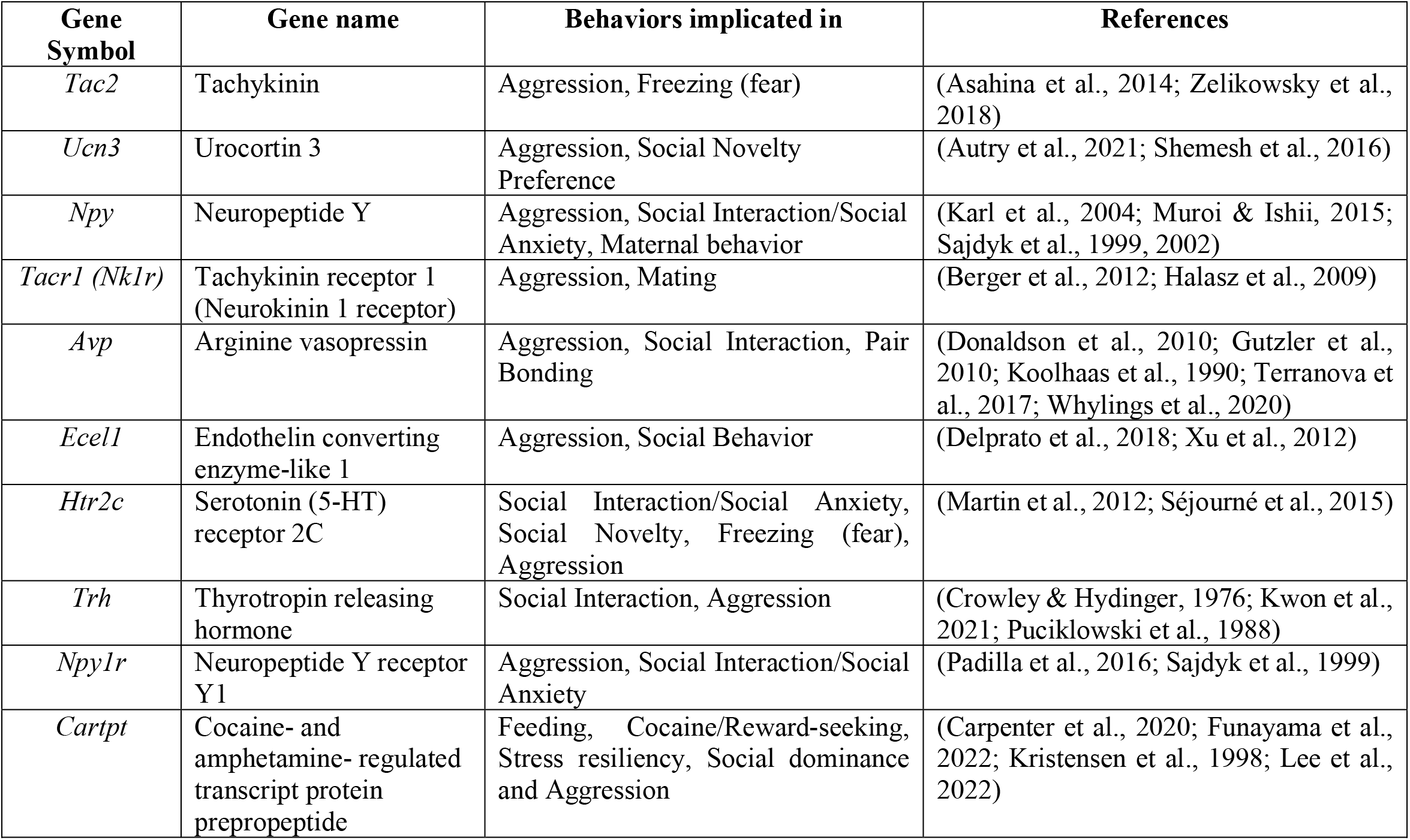
List of neuropeptide and receptor candidate genes expressed in the MeA and previously implicated in social or other behaviors.

**Figure 6:**
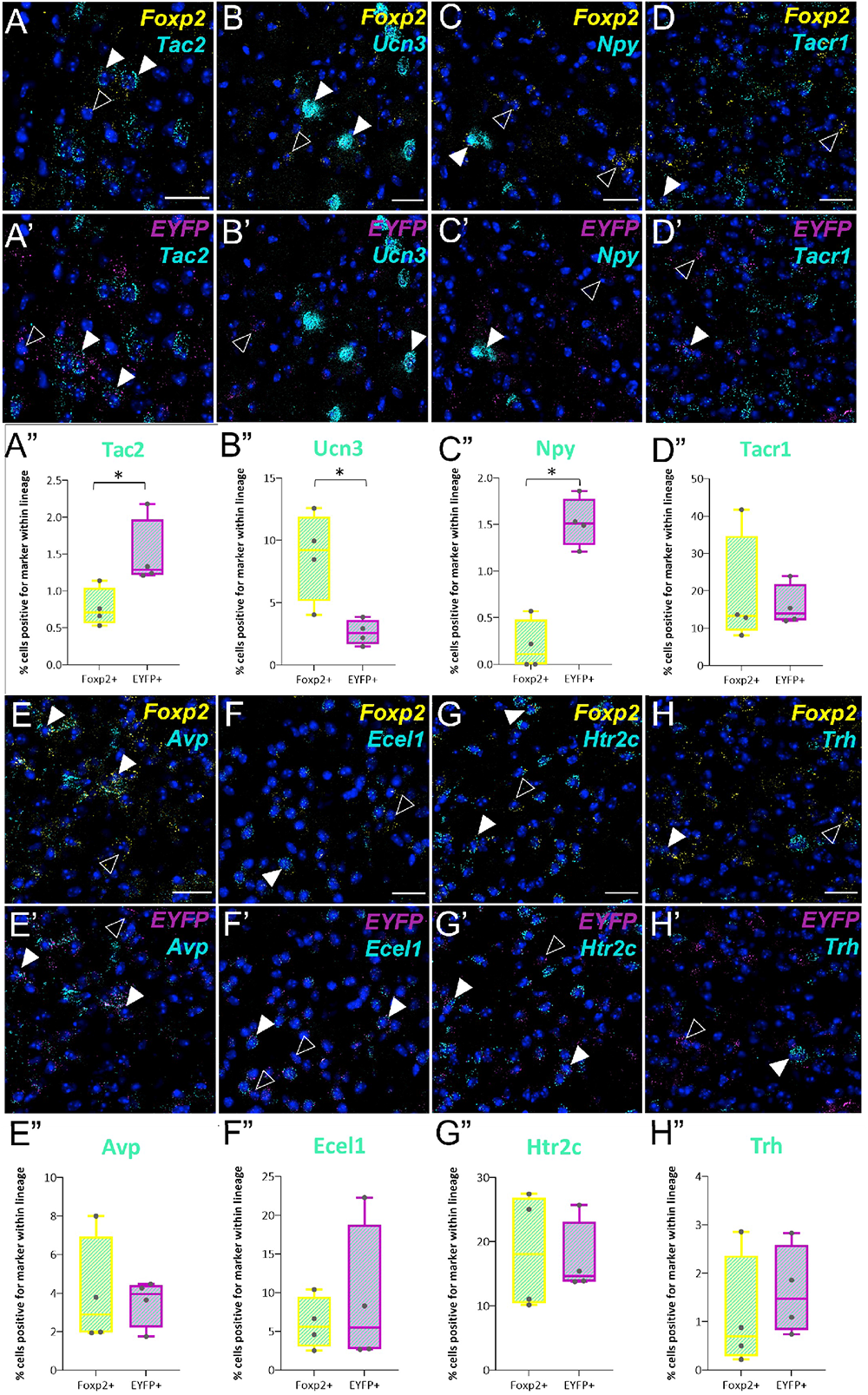
Fluorescent *in situ* hybridization for neuropeptides and receptors in *Foxp2+* and *Dbx1-*lineage neurons in the MeA. (**A-A”**) Expression and quantification of *Tac2* mRNA (cyan)-expressing neurons within *Foxp2* mRNA*+* (yellow) and *Dbx1-*lineage mRNA+ (magenta) cells as shown in coronal sections from the MeA. Box-and-whisker plots show percentage of co-expression of *Tac2* mRNA within each lineage (**A”**). (**B-H’’**) Expression and quantification of the following other mRNAs: *Ucn3* (**B-B”**), *Npy* (**C-C”**), *Tacr1* (**D-D”**), *Avp* (**E-E”**), *Ecel1* (**F-F”**), *Htr2c* (**G-G”**), *Trh* (**H-H”**). Solid white arrowheads indicate neurons double positive for mRNAs of candidate gene and *Foxp2* or *EYFP*; empty arrowheads indicate neurons positive for *Foxp2* or *EYFP* but not the candidate gene mRNA. Scale bars: 25 µm for all images. * FDR-adjusted *p* value < 0.05. All other comparisons not significant (*p*>0.05). n = 4 mice (2-4 sections per mouse, bilateral counts).

### 3.4 Sex differences in inhibitory and excitatory input to MeA *Foxp2*- and *Dbx1*-lineages

In addition to molecular differences described above and in Lischinsky et al. (2017) between MeA *Foxp2-* and *Dbx1-*lineage neurons, our previous studies revealed lineage differences in intrinsic biophysical properties (Matos et al., 2020). Prior studies from others revealed male/female differences in total inputs to the MeA, with males displaying more excitatory input (Billing et al., 2020; Cooke & Woolley, 2005). However, the MeA neuronal subtype target of these inputs remained unknown. Therefore, we next conducted patch clamp electrophysiology and measured spontaneous postsynaptic currents (sPSCs), a measure of homeostatic activity, in YFP+ MeA neurons in *Foxp2^cre^;YFP* and *Dbx1^cre^;YFP* mice (**Figure 7A,B**). Recordings in artificial cerebrospinal fluid (ACSF) alone, measured all spontaneous currents. To distinguish the contributions of different excitatory and inhibitory inputs, we blocked GABAA receptor-mediated (GABAAR) currents with PTX alone or both GABAA receptor-mediated currents and NMDA receptor-mediated (NMDAR) currents with PTX and AP5 (**Figure 7C**). We found a significantly higher frequency of total sPSCs (**Figure 7D**) in male *Dbx1*-lineage neurons compared to male *Foxp2*-lineage neurons. We additionally observed that while there was no sex difference within the *Dbx1*-lineage, *Foxp2*-lineage neurons in females displayed a significantly higher event frequency than those in males (**Figure 7D**). Blocking GABAAR currents with PTX alone (**Figure 7E**) or blocking GABAAR and NMDAR currents with PTX and AP5 (**Figure 7F**) eliminated the sex differences observed in frequency of total sPSCs in *Foxp2*-lineage neurons. However, *Foxp2*-lineage neurons in both conditions continued to display a lower frequency of events than *Dbx1*-lineage neurons, but only in males. This suggests that *Foxp2*-lineage neurons receive lower excitatory input than *Dbx1*-lineage neurons in males, a finding consistent with our previous observations (Lischinsky et al., 2017). In addition, *Foxp2*-lineage neurons receive overall more inhibitory input in females than males as those were significantly reduced in females when PTX was introduced, abolishing the sex difference within the *Foxp2-*lineage.

**Figure 7:**
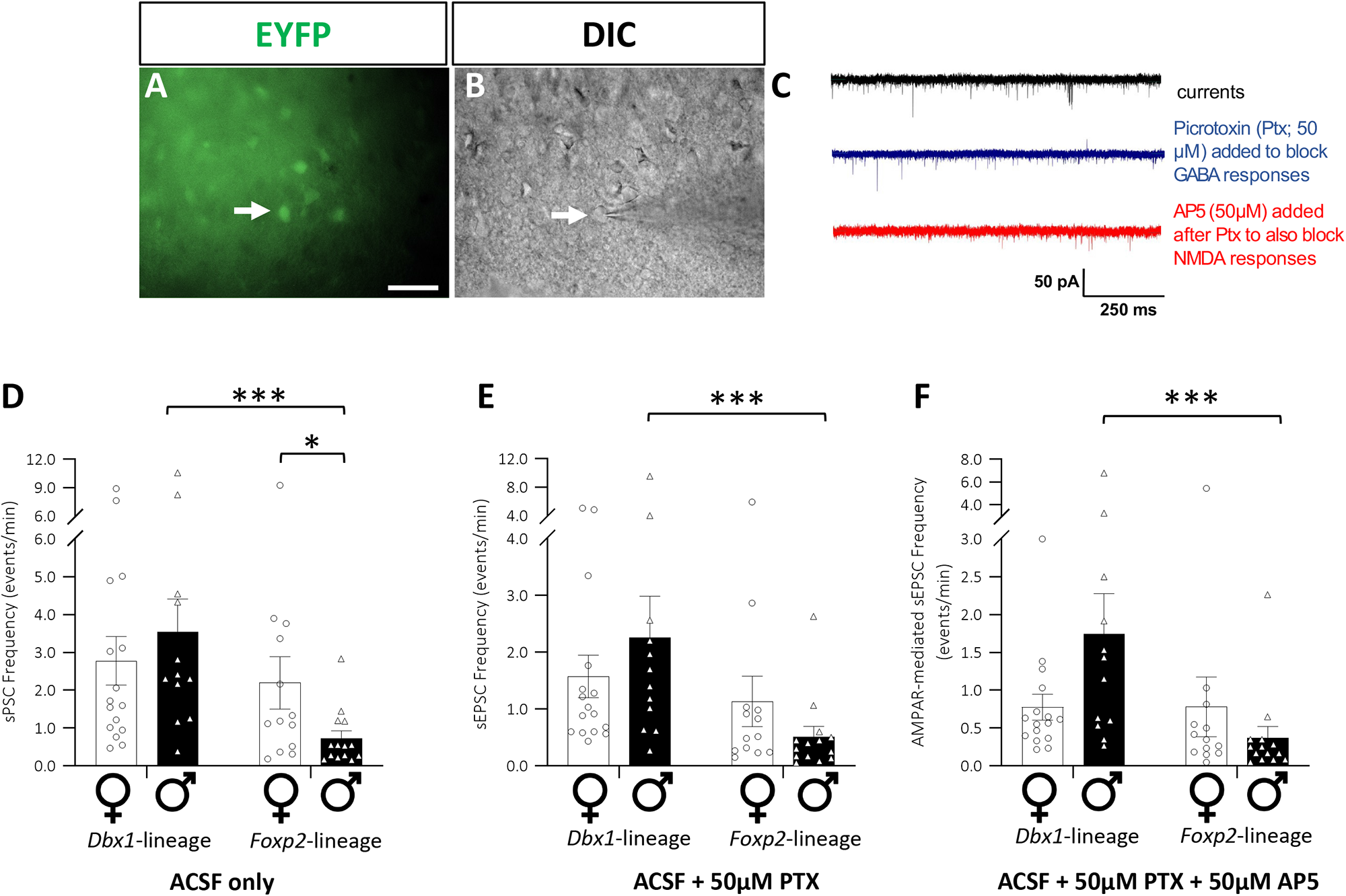
Sex- and lineage-differences in frequency of spontaneous postsynaptic currents (sPSCs) (**A**) Representative epifluorescence and (**B**) DIC images of the MeA showing a recombined EYFP+ neuron (arrow) targeted for *ex vivo* patch electrophysiology. (**C**) Representative current traces in whole-cell voltage clamp mode: in ACSF only (black, top), with 50 µM picrotoxin added (blue, middle) and with 50 µM AP5 added after picrotoxin (red, bottom). (**D-F**) Quantification of frequency (events per minute plotted as mean ± s.e.m) of sPSCs in *Dbx1*-(left) or *Foxp2*-lineage (right) neurons in female (circles, white bar) or male (triangles, black bar) mice with (**D**) bath ACSF only (all sPSCs), (**E**) bath-added 50 µM picrotoxin (sEPSCs only) or (**F**) 50 µM AP5 bath-added after picrotoxin (AMPAR-mediated sEPSCs only). * *p* < 0.05, *** *p* < 0.001, all other pairwise comparisons not significant, α=0.05. (n=12-16 neurons from 5-7 mice per group). Scale bar: 50 µm in (**A, B**)

As we observed sex differences in recordings with no blockers within MeA *Foxp2*-lineage neurons (**Figure 7D**), we next wanted to assess whether there were corresponding differences in the expression of either excitatory or inhibitory synaptic markers on MeA *Foxp2*-lineage neurons. To accomplish this, we assessed expression of excitatory pre- and postsynaptic markers, PSD-95 and VGLUT2 (**Figure 8**) and inhibitory pre- and postsynaptic markers Gephyrin and VGAT, (**Figure 9**) in *Foxp2^cre^;RYFP* mice by immunohistochemistry. We quantified both the proportion of *Foxp2*-lineage (YFP+) neurons receiving excitatory and inhibitory inputs as well as the number of colocalized pre- and postsynaptic excitatory or inhibitory puncta on YFP+ neurons. We established a threshold of 5 or more puncta on a YFP+ neuron to count it as positive for the corresponding marker. We found that a subset of *Foxp2*- lineage neurons in both males and females receive putative direct excitatory (**Figure 8A-E’**) or inhibitory (**Figure 9A-E’**) input. However, we found no differences in the proportion of neurons receiving excitatory (**Figure 8F**) or inhibitory (**Figure 9F**) input in males compared to females, nor in the number of puncta on each YFP+ neuron (data not shown).

**Figure 8:**
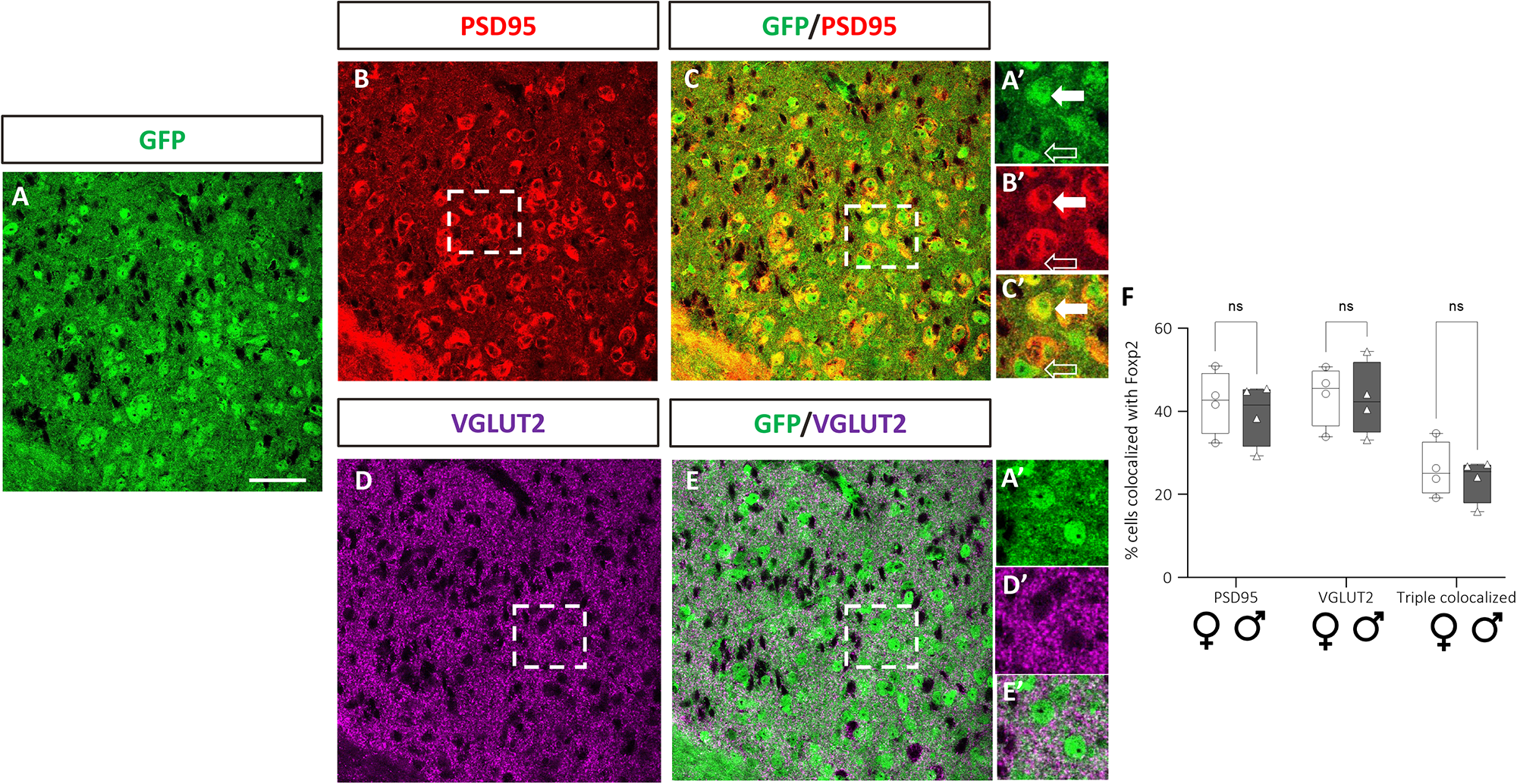
MeA *Foxp2*-lineage cells receive excitatory inputs, with no sex differences. (**A**) MeA *Foxp2*-lineage cells identified by GFP immunofluorescence (green) in coronal sections from *Foxp2^cre^;RYFP* mice. (**B**) Immunofluorescence for the postsynaptic marker PSD95 (red) marks neurons receiving excitatory inputs. (**C**) Co-immunofluorescence for GFP (green) and PSD95 (red). Boxed area in (**B, C**) shown at higher magnification in (**A’, B’, C’**) highlights a cell co-expressing PSD95 and GFP (arrow) and a cell with minimal co-expression (open arrow). (**D**) VGLUT2 immunofluorescence (red) marks presynaptic excitatory input. (**E**) Co-immunofluorescence for GFP (green) and VGLUT2 (magenta) reveals extensive excitatory inputs within the region of *Foxp2*-lineage cells. Boxed area in (**D, E**) shown at higher magnification in (**A’, D’, E’**) highlights *Foxp2*-lineage cells (green) receiving numerous VGLUT2+ inputs (magenta). (**F**) Box-and-whisker plot quantification of the percentage of *Foxp2*-lineage cells colocalized with PSD95 puncta, VGLUT2 puncta or both reveals no significant differences between females (circles, white bar) and males (triangles, grey bar). Scale bar in **A** equals 100 µm (also for **B-E**). Scale bar in **A’-E’** equals 20 µm.

**Figure 9:**
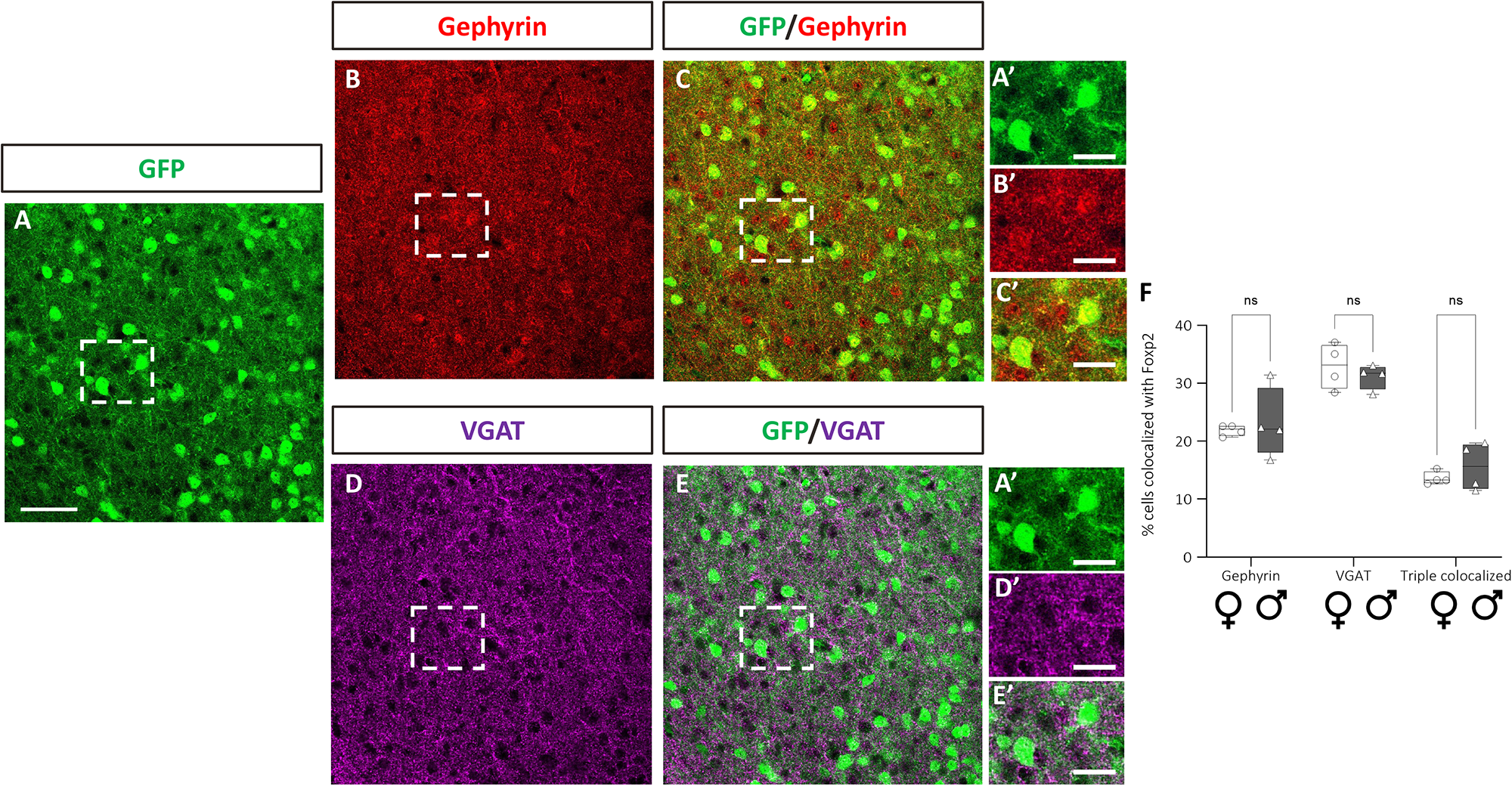
MeA *Foxp2*-lineage cells receive inhibitory inputs, with no sex differences. (**A**) MeA *Foxp2*-lineage cells identified by GFP immunofluorescence (green) in coronal sections from *Foxp2^cre^;RYFP* mice. (**B**) Immunofluorescence for the postsynaptic marker Gephyrin (red) marks neurons receiving inhibitory inputs. (**C**) Co-immunofluorescence for GFP (green) and Gephyrin (red) reveals many Gephyrin expressing *Foxp2*-lineage cells. Boxed area in (**B, C**) shown at higher magnification in (**A’, B’, C’**) highlights *Foxp2*-lineage cells expressing Gephyrin. (**D**) VGAT immunofluorescence (red) marks presynaptic inhibitory input. (**E**) Co-immunofluorescence for GFP (green) and VGAT (magenta) reveals extensive inhibitory inputs in the region of *Foxp2*-lineage cells. Boxed area in (**D, E**) shown at higher magnification in (**A’, D’, E’**) highlights *Foxp2*-lineage cells (green) receiving numerous VGAT inputs (magenta). (**F**) Box-and-whisker plot quantification of the percentage of *Foxp2*-lineage cells colocalized with puncta of each inhibitory synaptic marker reveals no significant differences between females (circles, white bar) and males (triangles, grey bar). Scale bar in **A** equals 100 µm (also for **B-E**). Scale bar in **A’-F**’ equals 20 µm.

## 4. Discussion

### 4.1 Summary of findings

Using a combination of gene expression analyses and virus-based circuit mapping approaches, we studied the molecular diversity and patterns of connectivity of neurons that comprise the core structures, VNO, AOB and MeA, of the accessory olfactory system (AOS). We find that two transcription factor-expressing populations, Foxp2+ neurons and neurons derived from the *Dbx1*-lineage define molecularly distinct neuronal populations across brain nuclei that comprise the AOS. We further find that Foxp2+ neurons in the AOB comprise the overwhelming majority of outputs to the MeA. Interestingly, we further find sex differences in the frequency of inputs to MeA *Foxp2*-lineage neurons, and within males only, lineage differences in the frequency of inputs. Thus, our findings suggest that subgroups of neurons identified by current or prior expression of select transcription factors may define distinct subcircuits with in the AOS and uncover a sexual dimorphism in their connectivity.

### 4.2. Lineage diversity across an interconnected circuit

The AOS is primarily dedicated to the processing of innate behaviors such as mating, aggression and predator avoidance. These behaviors are considered ‘hardwired’; meaning that they manifest without prior training. Although shaped by hormonal influences, the patterns of wiring of these circuits are likely in large part pre-determined by developmental genetic programs. However, these genetic programs remain unknown. Our previous studies, and the studies of others linking MeA embryonic development to neuronal diversity (Carney et al., 2010; Lischinsky et al., 2017) revealed that embryonic expression of the transcription factors Otp, Foxp2 and Dbx1 define separate populations of neural progenitors thatlater give rise to non-overlapping populations of either excitatory (Otp) or inhibitory (Foxp2, Dbx1) MeA output neurons (Lischinsky et al., 2017). Across the developing nervous system, expression of discrete subclasses of transcription factors in neural progenitors directs the emergence of later neuronal subtype identity (Aydin et al., 2019; Heavner et al., 2020; Hörmann et al., 2020; Sagner et al., 2021). The early endowment of MeA neuronal diversity as defined by transcription factor expression (Otp, Foxp2, Dbx1), suggested a potential molecular code for how the MeA is assembled. Here, our molecular analysis revealed that this molecular coding may be at least partially conserved in the VNO and AOB which lie one and two nodes upstream of the MeA, respectively. We found that in addition to the MeA, *Dbx1*-lineage neurons mark subsets of sensory neurons in the VNO and output neurons in the AOB. In the VNO, *Dbx1*-lineage neurons comprise ∼7% of the entire sensory neuron population across both apical and basal layers. Concurrent work has revealed that Dbx1 is expressed as early as E12.5 in the developing VNO (Causeret et al., 2022). Thus, similar to other regions of the nervous system, Dbx1 marks early developing progenitors in the VNO. As the VNO is the first site of sensory processing in the AOS, it is interesting to speculate that *Dbx1*-lineage sensory neurons may express specific subclasses of olfactory receptors implicated in select innate behaviors. If this is turns out to be the case, this also raises the interesting question of whether neurons of the same transcription factor identity/lineage directly connect with each other to form a transcription factor labeled line for the processing of select olfactory cues. However, the case for a such labeled-line circuit is not as strong for Foxp2. In contrast to the *Dbx1*-lineage, Foxp2 marks a much larger group of AOB neurons, comprising more than 2/3^rds^ of the output neurons projecting directly to the MeA. The lower number of *Dbx1*-lineage neurons in the AOB precluded our ability to trace *Dbx1*-lineage AOB projections to their final destinations using anterograde viral tracing. However, it would be reasonable to assume that AOB *Dbx1*-lineage neurons in part comprise the MeA-projecting Foxp2-negative population. Regardless, our findings here reveal that beyond the MeA, expression of the same transcription factors define regions of a known interconnected circuit.

### 4.3. MeA Neuropeptide Expression

Our previous studies revealed that *Foxp2*- and *Dbx1*-lineage neurons in the MeA express cohorts of neurohormones and ion channels in a lineage-specific manner. The neurohormones Aromatase and ER-α, and the action potential regulating ion channels *Kir5.1, Kir6.1, KChip4.1, Cav1.2* and *Kv7.1* are expressed more in *Dbx1*-lineage neurons, with the ion channel *Kir2.1* expressed more in the *Foxp2*-lineage (Lischinsky et al., 2017; Matos et al., 2020). Here, we extended this prior knowledge by assessing expression of neuropeptides known to have a function in MeA-regulated behaviors such as feeding, aggression, and mating. We show that expression of CARTPT, *Tac2* and *Npy* are enriched in the *Dbxl*-lineage while *Ucn3* is expressed in more *Foxp2*+ than *Dbx1-* lineage cells. Of these, the most striking pattern was with CARTPT, whose expression pattern strikingly mimics the distribution of *Dbx1*-lineage cells. As CARTPT antibody marks both cell bodies and CARTPT+ fibers, it was difficult to discern if *Dbx1*-lineage neurons are producing CARTPT or receiving dense CARTPT+ input. Regardless, the strong expression overlap implicates *Dbx1*-lineage neurons in aspects of feeding, homeostasis and/or reward. In addition, our multiplexed RNAscope *in situ* hybridization analyses revealed a higher expression of *Tac2* and *Npy* in *Dbx1*-lineage cells further implicating *Dbx1*-lineage neurons in aggression, social novelty or sexual arousal, which are known functions of these neuropeptides (Asahina et al., 2014; Karl et al., 2004; Zelikowsky et al., 2018). In contrast, of the 10 neuropeptides explored, we found only *Ucn3* to be enriched in the *Foxp2*+ population. Ucn3 is implicated in social novelty preference, infant-directed aggression and feeding (Autry et al., 2021; Shemesh et al., 2016; Stengel & Taché, 2014), suggesting a role for Foxp2+ neurons in these behaviors and consistent with its known role in aggression (Herrero et al., 2021; Lischinsky et al., 2017, 2022). Aside from CARTPT, it is important to note that although lineage restricted, *Tac2, Npy* and *Ucn3* are only expressed in a small subset of each lineage. This, however, does not preclude a putative important lineage-specific role in behavior for the following reasons: First, our expression analysis was conducted on tissue from home-cage animals. As levels of neuropeptide expression are typically state-dependent, it is likely we are only observing the baseline of expression. Second, our prior studies of cFos activation patterns in the MeA revealed that even during robust behavioral tasks, cFos is only expressed in a subset of neurons within each lineage (Lischinsky et al., 2017). This indicates that perhaps only a handful of neurons within a given population needs to be active to be engaged in given behavior. Third, these (and other) neuropeptides may act in concert in overlapping or different populations to modulate behavior. While these lineage-specific expression patterns of neuropeptides importantly extend our knowledge of molecular diversity of MeA neurons, it remains to be determined which behaviors are regulated by each lineage, as there are many mechanisms beyond neuropeptide expression that influence behavior. These include, for examples, patterns and types of input/output connectivity and intrinsic neuronal excitability and other biophysical parameters. Future transcriptomic analysis of *Foxp2*- and *Dbx1*-lineage cells will provide a full picture of the molecular diversity of these populations.

### 4.2. Sex differences in connectivity

The MeA has been long recognized as a highly sexually dimorphic brain region, with known differences in structural properties such as cell morphology, dendritic complexity, and cell size (Cooke et al., 2007; Cooke & Woolley, 2005; Hines et al., 1992). Our prior patch clamp electrophysiology studies further revealed sex differences in intrinsic biophysical properties, including action potential firing dynamics in both *Foxp2*- and *Dbx1*-lineages (Matos et al., 2020). Moreover, recent gene expression and single-cell RNA-seq transcriptomic studies revealed sex differences at the molecular level in the MeA (Chen et al., 2019) . Interestingly, the sex differences in gene expression in the MeA were most prominent in inhibitory GABAergic neurons as opposed to the excitatory glutamatergic population.

While our previous work revealed that *Dbx1*-lineage neurons receive greater excitatory input than *Foxp2*-lineage neurons (Lischinsky et al., 2017), in that prior study we did not explore whether there were sex differences in inputs to these two GABAergic output populations. To examine this, here we took both an electrophysiological approach by measuring the frequency of spontaneous inputs and an immunohistochemical approach to assess the status of synaptic connectivity using well characterized markers of excitatory and inhibitory synapses. Via patch clamp analyses, we uncovered sex differences in total inputs to *Foxp2*-lineage neurons, but not to *Dbx1*-lineage neurons. We found *Foxp2*-lineage neurons in females have more spontaneous responses than in males. These differences were no longer significantly different when GABAAR currents alone were blocked or when both GABAAR and NMDAR currents were blocked simultaneously. There are several possible mechanisms that can account for these observations in the *Foxp2*-lineage. First, the number of excitatory and/or inhibitory inputs to *Foxp2*-lineage cells maybe sexually dimorphic. However, our immunohistochemical analysis of synaptic markers suggests that this may not be the case. Although we cannot rule out that more direct analysis of synapses via ultrastructural analyses would uncover differences, our data suggest other mechanisms maybe occurring, such as sex differences in presynaptic firing rate and/or number of presynaptic neurotransmitter release events or postsynaptic membrane excitability via differences in ion channel expression. This finding could explain fewer postsynaptic currents that we observe in this study. Further clues to the underlying mechanism maybe inferred from recent MeA RNA-seq studies (Chen et al., 2019), which revealed major transcriptomic sex differences in MeA GABAergic neurons in genes implicated in synaptic function and communication. Although we do not know which GABAergic subpopulations give rise to these transcriptomic differences, our findings provide an entry point to link sex differences in gene expression with sex differences in synaptic input that we observe in MeA *Foxp2*-lineage neurons.

Interestingly, prior electrophysiological studies revealed that the MeA receives more excitatory input in males than in females (Cooke & Woolley, 2005). A more recent anatomical study showed that MeA Aromatase+ neurons in males receive more inputs from the AOB than females (Dwyer et al., 2022). While in potential contrast to our findings of greater synaptic input in females, it is likely that the male/female pattern of inputs to the MeA varies from population to population; with some neuronal subtypes receiving more (or stronger) inputs in males and others in females. In addition to the *Foxp2*-and *Dbx1-*lineage neurons, the MeA is populated by a vast array of interneurons, excitatory (Otp+ neurons) output neurons (Chen et al., 2019; Lischinsky et al., 2017) and perhaps non-*Foxp2*- and *Dbx1*-lineage inhibitory output neurons. The MeA receives strong input from not only the AOB, but from the posterior amygdala, cortical amygdala, BNST as well as lesser input from other brain regions (Lischinsky et al., 2022). Thus, there are likely sex differences not only based on which MeA population is innervated but also by source of input. Regardless of the underlying mechanism, our study introduces an additional layer of refinement in the analysis of physiological inputs to the MeA viz. developmental transcription factor-defined subpopulation heterogeneity.

Although we found robust differences across sex in synaptic input as uncovered by patch clamp electrophysiology, it is important to note that in our analysis we did not segregate females based on estrus cycle. Prior work in the MeA (Dalpian et al., 2019) and the hypothalamus (Dias et al., 2021; Yin et al., 2022), revealed estrus state-dependent changes in the strength of neuronal connectivity. Moreover, hormones greatly shape brain development at several levels. In addition to the estrus state-dependent short-term hormonal cycling which transiently affects circuitry, developmental hormonal surges also have an impact on how these circuits are initially wired together (Simerly, 2003). As the MeA plays a central role in regulating sex-specific innate behaviors such aggression and mating, there may also be non-hormonally driven genetic programs that are involved in the establishment of male and female differences in wiring patterns during early postnatal development. Exploration of both these intrinsic and extrinsic influences on induction and maintenance of sexually dimorphic patterns of MeA connectivity and ultimately behavior will be an interesting area of investigation.

## Acknowledgements

We kindly acknowledge intellectual input from members of the Corbin lab, present and past, and constructive feedback from the Haydar lab in the Center for Neuroscience Research at Children’s National.

## Author Contributions

NP, HM and SS were involved in the design, implementation, data generation, analysis and interpretation and oversaw trainees and staff in this project. NP with JGC wrote and edited the manuscript. NP generated the data in Figure 6. HM generated the data in Figure 7. SS generated the data in Figures 3, 8 & 9. LT under the direction of HM generated the data in Figure 4. TT under the direction of KS generated the data in Figure 1. JL, TH and NP generated data in Figure 2. SE and JL generated the data in Figure 5. AA, IA, NC, MG, DH-P, WM assisted in technical aspects of data generation and analysis. MH, ST, SY, YIK, NP generated probe lists for Table 1. KH-T oversaw and worked with ST, SY and YI. KJ co-mentored HM with JGC. JGC oversaw all aspects of the project from conception to execution.

## Data Availability Statement

The data that support the findings of this study are available from the corresponding author upon reasonable request.

